# Antibacterial T6SS effectors with a VRR-Nuc domain induce target cell death via DNA Double-Strand Breaks

**DOI:** 10.1101/2021.12.26.474169

**Authors:** Julia Takuno Hespanhol, Daniel Enrique Sanchez Limache, Gianlucca Gonçalves Nicastro, Liam Mead, Edgar Enrique Llontop, Gustavo Chagas Santos, Chuck Shaker Farah, Robson Francisco de Souza, Rodrigo da Silva Galhardo, Andrew Lovering, Ethel Bayer Santos

**Affiliations:** Departamento de Microbiologia, Instituto de Ciências Biomédicas, Universidade de São Paulo, São Paulo 05508-900, Brazil; Department of Biosciences, University of Birmingham, Birmingham, UK; Departamento de Bioquímica, Instituto de Química, Universidade de São Paulo, São Paulo 05508-000, Brazil

**Keywords:** DUF3396, Effectors, SPI-22, T6SS, VRR-Nuc

## Abstract

The T6SS (Type VI secretion System) secretes antibacterial effectors into target competitors. *Salmonella* spp. encode five phylogenetically distinct T6SSs. Here we characterize the function of the SPI-22 T6SS of *S. bongori*, showing that it has antibacterial activity. We identify a group of antibacterial T6SS effectors (TseV1-4) containing an N-terminal PAAR-like domain and a C-terminal VRR-Nuc domain encoded next to cognate immunity proteins that contain the DUF3396 domain (TsiV1-4). TseV2 and TseV3 are toxic when expressed in *Escherichia coli* and bacterial competition assays confirm that TseV2 and TseV3 are secreted by the SPI-22 T6SS. Phylogenetic analysis reveals that TseV1-4 are evolutionarily related to enzymes involved in DNA repair. TseV2 and TseV3 maintained the ability to bind DNA, but instead cause specific DNA double-strand breaks and induce the SOS response in target cells. The crystal structure of the TseV3:TsiV3 complex reveals that the immunity protein likely blocks the effector interaction with the DNA substrate. These results expand our knowledge on the function of *Salmonella* pathogenicity islands, the evolution of toxins used in biological conflicts, and the endogenous mechanism regulating the activity of these toxins.

## Introduction

Bacteria use a series of antagonistic mechanisms to counteract competitors. These processes either require physical contact between attacker and target cells or function in a contact-independent manner via soluble molecules secreted into the medium (Peterson *et al*, 2020). The type VI secretion system (T6SS) is a multi-protein contractile nanomachine evolutionarily related to bacteriophages (Leiman *et al*, 2009). This system is widespread in Gram-negative bacteria and secretes toxic effectors into target cells in a contact-dependent manner (Coulthurst, 2019). The T6SS is composed of three major complexes: the membrane complex, the baseplate and the tail (Nguyen *et al*, 2018). The tail has a spear-like shape and is propelled against target cells upon a contraction event (Wang *et al*, 2017; Salih *et al*, 2018). The tail tube is composed of hexameric rings of Hcp (hemolysin co-regulated protein) capped with a spike composed of a trimer of VgrG (valine-glycine repeat protein G) and a PAAR protein (proline-alanine-alanine-arginine repeats) (Mougous *et al*, 2006; Shneider *et al*, 2013; Renault *et al*, 2018). The effectors secreted via T6SSs associate with Hcp, VgrG or PAAR either directly or indirectly via adaptor proteins (cargo effectors). In addition, so-called evolved effectors are fused to the C-terminus of Hcp, VgrG or PAAR (Cianfanelli *et al*, 2016; Jana & Salomon, 2019). Several isoforms of VgrG, Hcp and PAAR proteins can be encoded in the same bacterial genome, usually outside of the T6SS structural gene cluster (and are thus named orphan proteins). These Hcp, VgrG and PAAR proteins can assemble in different combinations to secrete specific subsets of effectors (Hachani *et al*, 2014; Bondage *et al*, 2016).

T6SSs effectors can target eukaryotic cells, prokaryotic cells or contribute to the acquisition of micronutrients (Coulthurst, 2019). The variety of targets is related to the diversity of biochemical activities of T6SS effectors, which can be nucleases, peptidoglycan hydrolases, lipases, NADases, pore-forming proteins or enzymes that post-translationally modify target proteins (Jurėnas & Journet, 2021). Antibacterial effectors with nuclease activity are among the most potent weapons used by an attacker to intoxicate target cells. Several T6SS effectors with nuclease activity have been reported including, *Dickeya dadantii* RhsA-CT and RhsB-CT (Koskiniemi *et al*, 2013); *Agrobacterium tumefaciens* Tde1 and Tde2 (Ma *et al*, 2014); *Pseudomonas aeruginosa* PA0099 (Hachani *et al*., 2014), TseT (Burkinshaw *et al*, 2018) and Tse7 (Pissaridou, 2018); *Serratia marcescens* Rhs2 (Alcoforado Diniz & Coulthurst, 2015); *Escherichia coli* Hcp-ET1, -ET3 and -ET4 (Ma *et al*, 2017a), and Rhs-CT3, -CT4, -CT5, -CT6, -CT7 and -CT8 (Ma *et al*, 2017b); *Acinetobacter baumannii* Rhs2-CT (Fitzsimons *et al*, 2018); *Vibrio parahaemolyticus* PoNe (Jana *et al*, 2019); *Aeromonas dhakensis* TseI (Pei *et al*, 2021); and *Burkholderia gladioli* TseTBg (Yadav *et al*, 2021).

The majority of the nuclease domains mentioned above have been previously predicted by a seminal *in silico* study using comparative genomics (Zhang *et al*, 2012). Among those characterized are Ntox15 (PF15604) (Ma et al., 2014); Ntox30 (PF15532), Ntox34 (PF15606) and Ntox44 (PF15607) (Ma *et al*., 2017a); Tox-REase-1 (Jana et al., 2019); Tox-REase-3 (PF15647) (Ma *et al*., 2017a); Tox-REase-5 (PF15648) (Burkinshaw *et al*., 2018; Yadav *et al*., 2021); Tox-GHH2 (PF15635) (Hachani *et al*., 2014; Pissaridou, 2018); HNH (PF01844) (Koskiniemi *et al*., 2013; Alcoforado Diniz & Coulthurst, 2015; Ma *et al*., 2017b); Tox-JAB-2 (Ma *et al*., 2017a); AHH (PF14412) (Ma *et al*., 2017a; Fitzsimons *et al*., 2018); and Tox-HNH-EHHH (PF15657) (Pei *et al*., 2021).

In *Salmonella* species, T6SSs are encoded in five distinct *Salmonella* pathogenicity islands (SPI-6, SPI-19, SPI-20, SPI-21 and SPI-22) acquired by different horizontal gene transfer events (Blondel *et al*, 2009; Bao *et al*, 2019). The *S. enterica* serovar Typhimurium SPI-6 T6SS is involved in competition with the host microbiota and gut colonization (Pezoa *et al*, 2014; Brunet *et al*, 2015; Sana *et al*, 2016; Sibinelli-Sousa *et al*, 2020); whereas the SPI-19 T6SS of *S*. Gallinarium is involved in survival within macrophages (Blondel *et al*, 2013; Schroll *et al*, 2019). So far, only two T6SS effectors have been characterized in *Salmonella* spp., both targeting peptidoglycan: Tae4 (type VI amidase effector 4) is a gamma-glutamyl-D,L-endopeptidases that cleaves between D-*i*Glu^2^ and *m*DAP^3^ within the same peptide stem (Russell *et al*, 2012; Benz *et al*, 2013; Zhang *et al*, 2013); and Tlde1 (type VI L,D-transpeptidase effector 1), which exhibits both L,D-carboxypeptidase and L,D-transpeptidase D-amino acid exchange activity, cleaving between *m*DAP^3^ and D-Ala^4^ of the acceptor tetrapeptide stem or replacing the D-Ala^4^ by a noncanonical D-amino acid, respectively (Sibinelli-Sousa *et al*., 2020).

Herein we report the characterization of the SPI-22 T6SS of *S. bongori*, which displays antibacterial activity. We characterize a group of antibacterial effectors secreted by this system that contain a VRR-Nuc (virus-type replication-repair nuclease) domain (Kinch *et al*, 2005; Iyer *et al*, 2006), named type VI effector VRR-Nuc 1-4 (TseV1-4). These effectors are encoded next to DUF3396-containing proteins, which function as immunity proteins (TsiV1-4) that are specific to each effector. Phylogenetic analysis revealed that TseVs effectors form a clade together with other antibacterial effectors belonging to the PD-(D/E)xK phosphodiesterase superfamily. This toxic clade is phylogenetically related to enzymes containing the VRR-Nuc domain conventionally involved in DNA repair and metabolism. We confirm that TseV2 and TseV3 display DNAse activity, inducing the SOS response with resultant target cell death via DNA double-strand breaks. Our crystal structure of the TseV3:TsiV3 complex reveals that the immunity protein likely impairs effector toxicity by interacting with and occluding its DNA-binding site. Our results provide mechanistic knowledge about a new group of antibacterial toxins that co-opt the VRR-Nuc domain for a previously undescribed role in bacterial antagonism, and further reveal the mode of neutralization via specific immunity protein complexation.

## Results

### The SPI-22 T6SS of *S. bongori* has antibacterial activity

The SPI-22 T6SS of *S. bongori* is phylogenetically related to the HSI-III (Hcp secretion island III) T6SS of *Pseudomonas aeruginosa* (amino acid similarity ranging from 26-80%), and the CTS2 (*Citrobacter rodentium* T6SS cluster 2) of *C. rodentium* (amino acid similarity ranging from 63-93%) (Petty *et al*, 2010; Fookes *et al*, 2011) (Fig. 1A). Besides the structural T6SS components encoded within SPI-22, the genome of *S. bongori* NCTC 12419 encodes several orphan proteins comprising two VgrGs (SBG_2715, SBG_3770), four Hcps (SBG_0599, SBG_3120, SBG_3143, SBG_3925), three DUF4150/PAAR-like proteins (SBG_1846, SBG_2718, SBG_2955), two adaptors containing DUF2169 (SBG_1847, SBG_2721), and one adaptor with DUF1795 (SBG_3173) (Fig. 1B).

**Fig. 1.**
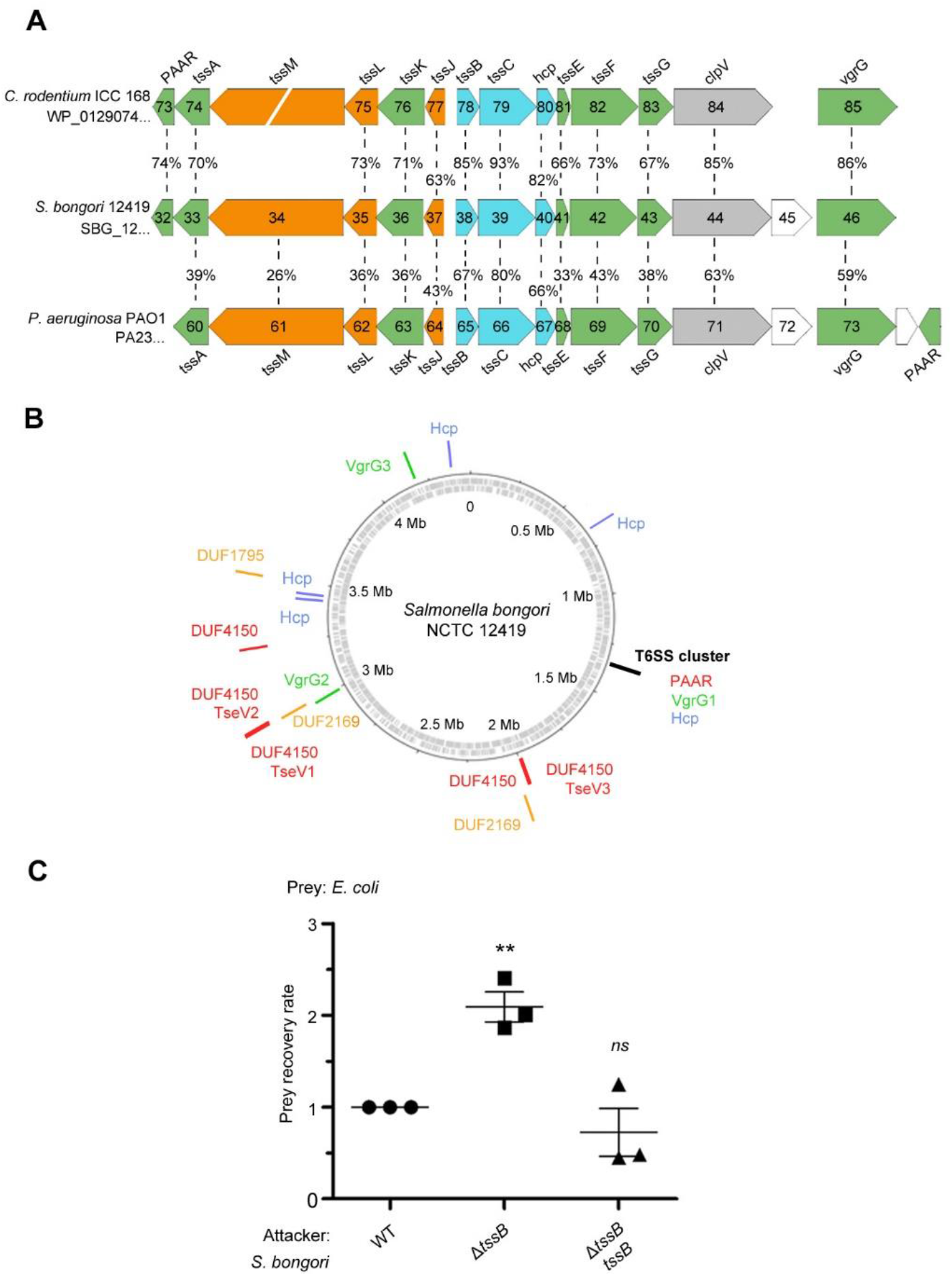
The *S. bongori* SPI-22 encodes an antibacterial T6SS. (A) Comparison between the SPI-22 T6SS of *S. bongori* with the systems of *C. rodentium* and *P. aeruginosa*. The T6SS proteins forming the three subcomplexes are in colors: membrane components (orange), sheath and inner tube (blue), and baseplate and spike components (green). (B) Representation of the circular genome of *S. bongori* with T6SS components highlighted: the structural cluster is marked by a black line; VgrG proteins are represented by green lines; Hcps are in blue; adaptor proteins are in orange; and PAAR or PAAR-like proteins are in red. TseV1, TseV2 and TseV3 fused to PAAR-like domain are also in red. (C) Bacterial competition assays between *S. bongori* WT, Δ*tssB* and Δ*tssB* complemented with pFPV25.1 *tssB* against *E. coli* in LB-agar incubated for 24 h. The prey recovery rate was calculated by dividing the colony-forming unit (CFU) counts of the output by the input. Data represent the mean ± SD of three independent experiments performed in duplicate and were analyzed through comparison with WT that were normalized to 1. One-way ANOVA followed by Dunnett’s multiple comparison test. **p < 0.01 and *ns* not significant.

To analyze whether *S. bongori* SPI-22 T6SS has *bona fide* antibacterial activity, we performed bacterial competition assays using the wild-type (WT) and T6SS null mutant (Δ*tssB/SBG_1238*) strains as attacker cells, and *Escherichia coli* K12 W3110 as prey. Results showed that the prey recovery rate was higher when co-incubation was performed with Δ*tssB* compared to the WT (Fig. 1C). In addition, competition with a Δ*tssB* strain complemented with a plasmid expressing TssB restored the WT phenotype (Fig. 1C). These results show that the SPI-22 T6SS of *S. bongori* is active in the conditions tested and contributes to interbacterial antagonism, thus priming investigation to further characterize this activity.

### TseV2 and TseV3 are antibacterial SPI-22 T6SS effectors

After verifying that the SPI-22 T6SS has antibacterial activity, we set out to identify the effectors contributing to the antagonistic effect. Initially, we performed *in silico* analysis using Bastion6 (Wang *et al*, 2018) to evaluate several candidates (10 genes up- and downstream of all T6SS components) (Fig. 1B) for their probability of being a T6SS effector (cutoff score ≥ 0.5) (data not shown). Two candidates called our attention: SBG_2718 (TseV1) and SBG_2723 (TseV2), which contain an N-terminal PAAR-like domain and a C-terminal VRR-Nuc domain (Fig. 2A) (Kinch *et al*., 2005; Iyer *et al*., 2006). Both putative effectors are encoded next to pairs of genes encoding DUF3396-containing proteins that resemble putative immunity proteins: SBG_2719/TsiV1.1 and SBG_2720/TsiV1.2, and SBG_2724/TsiV2.1 and SBG_2725/TsiV2.2 (Fig. 2A). Additional BLASTP searches in the genome of *S. bongori* identified two extra VRR-Nuc-containing proteins (SBG_1841/TseV3 and SBG_1828/TseV4), but only one of them encoding an N-terminal PAAR-like domain (SBG_1841). Similarly, SBG_1828 and SBG_1841 are encoded upstream of a DUF3396-containing protein (SBG_1829/TsiV4 and SBG_1842/TsiV3) (Fig. 2A).

**Fig. 2.**
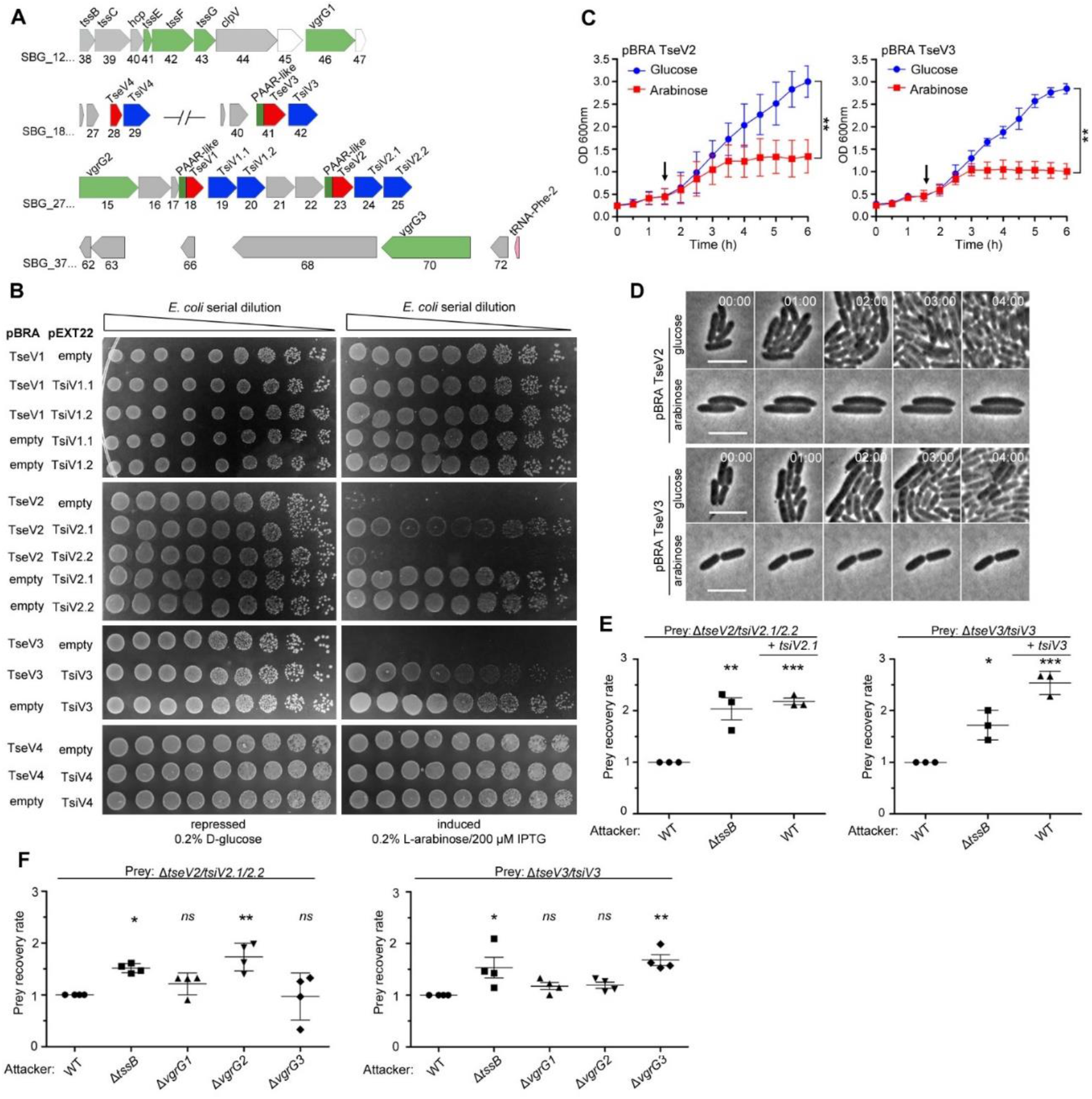
TseV2 and TseV3 are antibacterial SPI-22 T6SS effectors. (A) Scheme of the genomic region containing VgrGs and TseV/TsiV effector/immunity pairs. VRR-Nuc domain (red), PAAR-like domain (dark green), VgrG (light green), and DUF3396-containing immunities (blue). (B) *E. coli* toxicity assay. Serial dilutions of *E. coli* containing pBRA and pEXT22 constructs, as indicated, spotted onto LB agar plates, and grown for 20 h. Images are representative of three independent experiments. (C) Growth curve of *E. coli* harboring pBRA TseV2 or TseV3 before and after toxin induction by the addition of 0.2% L-arabinose (arrow). Results represent the mean ± SD of three independent experiments performed in duplicate. **p < 0.01 (Student’s *t* test). (D) Time-lapse microscopy of *E. coli* carrying either pBRA TseV2 or pBRA TseV3 grown on LB-agar pads containing either 0.2% D-glucose (repressed) or 0.2% L-arabinose (induced). Scale bar: 5 µm. Timestamps in hh:mm. (E) Bacterial competition assay using *S. bongori* WT, Δ*tssB* and Δ*tssB* complemented with pFPV25.1 *tssB* against *S. bongori* Δ*tseV2/tsiV2*.*1/tsiV2*.*2* or Δ*tseV3/tsiV3* complemented or not with pFPV25.1 *tsiV2*.*1* or pFPV25.1 *tsiV3*. Strains were co-incubated for 20 h (Δ*tseV2/tsiV2*.*1/tsiV2*.*2*) or 6 h (Δ*tseV3/tsiV3*) prior to measuring CFU counts. The prey recovery rate was calculated by dividing the CFU of the output by the input. Data represent the mean ± SD of three independent experiments performed in duplicate and were analyzed through comparison with WT that were normalized to 1. One-way ANOVA followed by Dunnett’s multiple comparison test. *p < 0.05, **p < 0.01 and ***p < 0.001. (F) Bacterial competition assay using *S. bongori* WT, Δ*tssB*, Δ*vgrG1*, Δ*vgrG2* or Δ*vgrG3* against *S. bongori* Δ*tseV2/tsiV2*.*1/tsiV2*.*2* or Δ*tseV3/tsiV3*. Strains were co-incubated for 20 h prior to measuring CFU counts. Prey recovery rate was calculated as in (E). Data represent the mean ± SD of four independent experiments performed in duplicate. One-way ANOVA followed by Dunnett’s multiple comparison test. *p < 0.05, **p < 0.01 and *ns* (not significant).

To analyze whether these proteins comprise four effector-immunity pairs, we cloned these genes into compatible vectors under the control of different promoters. To evaluate the toxicity of TseV1-4 upon expression in *E. coli*, the C-terminal regions of TseV1-3 and the full-length TseV4 were cloned into the pBRA vector under the control of the P_BAD_ promoter (inducible by L-arabinose and repressed by D-glucose). The putative immunity proteins were cloned into the pEXT22 vector under the control of the P_TAC_ promoter, which is inducible by isopropyl β-D-1-thiogalactopyranoside (IPTG). *E. coli* strains carrying different combinations of pBRA and pEXT22 were serially diluted and spotted onto LB agar plates containing either 0.2% D-glucose or 0.2% L-arabinose plus 200 μM IPTG (Fig. 2B). Results showed that TseV2 and TseV3 are toxic in the cytoplasm of *E. coli*, whereas TseV1 and TseV4 do not confer toxicity (Fig. 2B). Co-expression of TseV2 with either TsiV2.1 or TsiV2.2 revealed that only the first immunity protein neutralizes TseV2 toxicity (Fig. 2B). Similarly, the toxic effect of TseV3 can be neutralized by co-expression with TsiV3 (Fig. 2B). Co-expression of TseV2 and TseV3 with all combinations of immunity proteins (TsiV1.1, TsiV1.2, TsiV2.1, TsiV2.2, TsiV3 and TsiV4) revealed that the effectors are neutralized only by the specific cognate immunity protein (Fig. S1). The effect of TseV2 and TseV3 on cell growth was also analyzed in liquid media by measuring the OD_600ηm_ of *E. coli* carrying pBRA TseV2 or TseV3 (Fig. 2C). Under these conditions, bacteria grew normally in media containing D-glucose; but once L-arabinose was added, the culture stopped growing, and the OD_600ηm_ stabilized (Fig. 2C).

We performed time-lapse microscopy to evaluate growth and morphology of individual *E. coli* cells harboring pBRA TseV2 or TseV3. Bacteria grew normally when incubated in LB agar pads containing 0.2% D-glucose (repressed) over a time frame of 8 h (Fig. 2D, Movies S1, S3). However, in the presence of 0.2% L-arabinose (induced) bacteria did not grow and remained mostly morphologically unaltered – displaying a modest increase in cell length (Fig. 2D, Movies S2, S4).

To verify whether TseV2 and TseV3 are SPI-22 T6SS substrates, we performed bacterial competition assays using *S. bongori* WT and Δ*tssB* (attacker) versus *S. bongori* lacking either TsiV2.1/2.2 (Δ*tseV2/tsiV2*.*1/2*.*2)* or TsiV3 (Δ*tseV3/tsiV3)* as prey (Fig. 2E). Results demonstrated that the prey recovery rate was higher when prey cells were co-incubated with Δ*tssB* compared to WT (Fig. 2E). Complementation of preys with a plasmid encoding either TsiV2.1 or TsiV3 increased the prey recovery rate, showing that prey became immune to the TseV2 and TseV3-induced toxicity (Fig. 2E). These results confirm that TseV2 and TseV3 are antibacterial effectors secreted by the SPI-22 T6SS.

As TseV2 and TseV3 contain an N-terminal PAAR-like domain, which interacts with VgrG during T6SS assembly and effector secretion (Shneider *et al*., 2013), we decided to determine which of the three VgrG proteins encoded in *S. bongori* genome (Fig. 1B, 2A) were responsible for the secretion of TseV2 and TseV3. To shed light on this matter, we performed bacterial competition assays using *S. bongori* WT, Δ*tssB*, Δ*vgrG1* (SBG_1246), Δ*vgrG2* (SBG_2715) or Δ*vgrG3* (SBG_3770) (attacker) versus Δ*tseV2/tsiV2*.*1/2*.*2* or Δ*tseV3/tsiV3* (prey) (Fig. 2F). The prey recovery rate of Δ*tseV2/tsiV2*.*1/2*.*2* increased when this strain was co-incubated with Δ*vgrG2*, suggesting that VgrG2 is responsible for secreting TseV2 into target cells (Fig. 2F). Conversely, the prey recovery rate of Δ*tseV3/tsiV3* increased when this strain was co-incubated with Δ*vgrG3*, suggesting that VgrG3 is responsible for secreting TseV3 into target cells (Fig. 2F). VgrG2 and VgrG3 are 96.9% identical in their N-terminal region (VgrG2_1-565_ and VgrG3_1-545_) but display a distinct C-terminal domain with only 26% identity (VgrG2_566-709_ and VgrG3_546-728_) (Fig. S2), thus suggesting that this region is responsible for cargo selection (Liang *et al*, 2021). Together, these results show that each effector has its own mechanism of secretion, which is dependent on distinct VgrGs.

### VRR-Nuc-containing effectors are evolutionarily related to Holliday junction resolvases and enzymes involved in DNA repair

TseV2 and TseV3 contain a VRR-Nuc domain at their C-terminus, which was initially annotated as DUF994 (Kinch *et al*., 2005) and later renamed VRR-Nuc due to its association with enzymes linked to DNA metabolism (Iyer *et al*., 2006). VRR-Nuc-containing proteins are found in a wide range of organisms, including bacteria, bacteriophages, fungi, and eukaryotes (Iyer *et al*., 2006). Proteins containing this domain comprise a family (PF08774) belonging to the PD-(D/E)xK superfamily, which constitutes a large and functionally diverse group containing representatives involved in DNA replication (Holliday junction resolvases), restriction-modification, repair, and tRNA-intron splicing (Steczkiewicz *et al*, 2012). Members of this superfamily exhibit low sequence similarity but display a common fold in its enzymatic core (with α_1_β_1_β_2_β_3_α_2_β_4_ topology), which contains conserved residues (Asp, Glu, Lys) responsible for catalysis (Steczkiewicz *et al*., 2012).

To gain insight into the molecular function of TseV2 and TseV3 and understand their phylogenetic relationship, we used TseV1, TseV2 and TseV3 (TseV4 is 79.1% identical to TseV3 and was not used) amino acid sequences as queries in JackHMMER searches (Potter *et al*, 2018) for four iterations on the NCBI nr database (November 4th, 2021) to fetch a total of 2254 sequences with significant similarity (inclusion threshold ≤10^−9^ and reporting threshold ≤10^−6^). Additional JackHMMER searches were performed using selected VRR-Nuc-containing proteins as queries (Bce1019, PmgM, T1p21, KIAA1018, HP1472 and Plu1493) (Iyer *et al*., 2006), and recently reported *bona fide* or putative T6SS effectors that also belong to the PD-(D/E)xK superfamily: TseT (Burkinshaw *et al*., 2018); PoNe (Jana *et al*., 2019); IdrD-CT (Sirias *et al*, 2020); TseTBg (Yadav *et al*., 2021); Aave_0499 (Pei *et al*., 2021); and TseV^PA^ (Wang *et al*, 2021). A total of 39159 sequences were collected. For each JackHMMER dataset, we produced alignments with representatives from clusters formed by sequences displaying 80% coverage and 50-70% identity. These alignments were manually inspected, and divergent/truncated sequences were removed. We observed that the β_2_β_3_α_2_β_4_ region of the enzymatic core was more conserved so we used this region for a new multiple sequence alignment to build a phylogenetic tree using maximum likelihood (Fig. 3A).

**Fig. 3.**
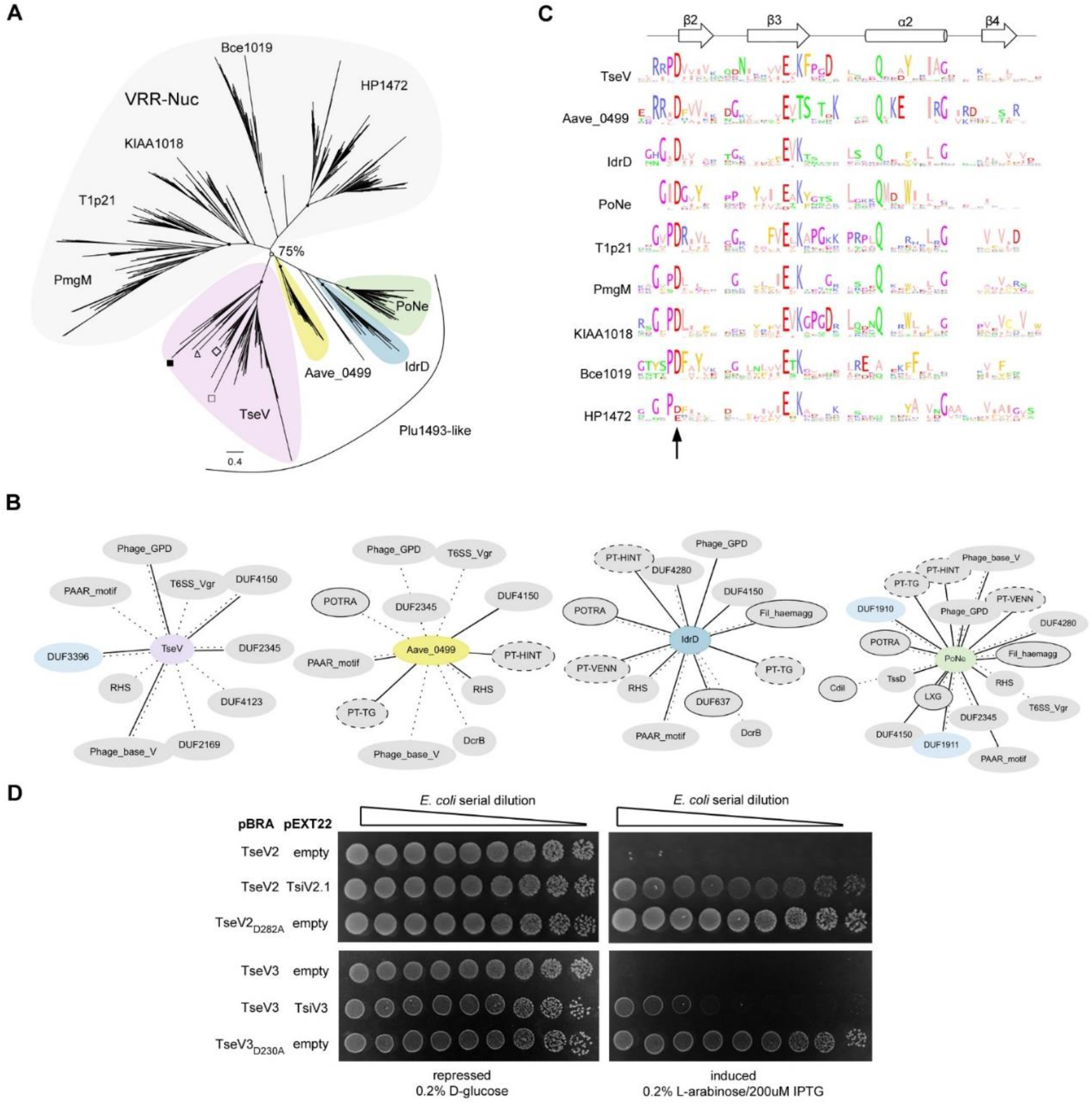
VRR-Nuc-containing effectors are evolutionarily related to enzymes involved in DNA metabolism. (A) Maximum likelihood phylogenetic tree of VRR-Nuc family members (Bce1019, PmgM, T1p21, KIAA1018, HP1472, Plu1493) (Iyer *et al*., 2006) and recently reported bona fide or putative T6SS effectors belonging to the PD-(D/E)xK superfamily (TseT, PoNe, IdrD-CT, TseTBg, Aave_0499, TseV^PA^). In the TseV clade (pink) the localization of TseV1 (□), TseV2 (Δ), TseV3 (■), and Plu1493 (◇) are marked. (B) Contextual network representation of domains and the genomic context of proteins belonging to Plu1493-like group (TseV, Aave_0499, IdrD, PoNe). Each circle represents a domain, which is either fused to (solid line) or encoded up- or downstream (dashed line) of the gene of interest (center). Borderless gray circles represent domains related to T6SS; bordered gray circles denote domains associated with a different bacterial secretion system; dashed nodes indicate pre-toxin domains; and light blue circles represent immunity proteins. (C) Sequence logo from the conserved β_2_β_3_α_2_β_4_ of the PD-(D/E)xK enzymatic core from all clades shown in (A). The arrow indicates conserved aspartic acid that was mutated in (D). (D) *E. coli* toxicity assay. Serial dilution of *E. coli* containing pBRA and pEXT22 constructs, as indicated, spotted onto LB agar plates, and grown for 20 h. Images are representative of three independent experiments.

The resulting tree is composed of 9 main clades, with 5 of these clades comprising PmgM, T1p21, KIAA1018, Bce1019 and HP1472 that reproduce the classification proposed by Iyer et al. (2006) in which each of these clades constitutes a subfamily of the VRR-Nuc family (Fig. 3A, grey; Table S1). Bce1019 subfamily contains the endonuclease I from Bacteriophage T7 (PDB 1M0D) (Hadden *et al*, 2002) and the transposon Tn7 encoded nuclease protein TnsA from *E. coli* (PDB 1F1Z) (Hickman *et al*, 2000) (PDB 1T0F) (Ronning *et al*, 2004). The PmgM subfamily contains a nuclease with the same name from phage P1 (Iyer *et al*., 2006). The T1p21 subfamily contains proteins encoded upstream of helicases (Iyer *et al*., 2006). The KIAA1018 group includes the human Fanconi anemia-associated nuclease 1 (FAN1) (PDB 4REA) (PDB 4RIA) and its bacterial homolog *Pa*FAN1 (PDB 4R89), which are involved in DNA repair (Gwon *et al*, 2014; Wang *et al*, 2014; Zhao *et al*, 2014). Curiously, antibacterial T6SS effectors formed 4 groups (Fig. 3A, colors) in which TseV2 and TseV3 clustered with Plu1493 (Iyer *et al*., 2006) and TseV^PA^ (Wang *et al*., 2021), whereas homologs of Aave_0499 (Pei *et al*., 2021), IdrD (Sirias *et al*., 2020) and PoNe (Jana *et al*., 2019) formed separated clades (Fig. 3A, colors; Table S1). These results indicate that TseV proteins are members of the Plu1493 subfamily (Iyer *et al*., 2006). Conversely, homologs of TseT were too divergent to be grouped in the phylogenetic tree and impaired its reproducibility, thus indicating that they probably have a distinct evolutionary origin (Fig. S3; Table S1).

All T6SS effectors (TseVs, Aave_0499, IdrD and PoNe), except for TseT homologs, formed a clade with a bootstrap value higher than 75% (Fig. 3A, colors). The genomic context of TseV/Plu1493 homologs is different from the other VRR-Nuc family members (Table S2). While most of VRR-Nuc members (PmgM, T1p21, KIAA1018, Bce1019 and HP1472) are encoded next to genes involved in DNA metabolism, the gene neighborhood of antibacterial T6SS effectors (TseVs, Aave_0499, IdrD and PoNe) is enriched in proteins encoding components of the T6SS apparatus, adaptors, and immunity proteins (Fig. 3B; Table S2). In addition, we observed proteins containing domains of other secretion systems involved in biological conflicts, such as CdiB and POTRA (T5SS) and LXG (T7SS) domains (Fig. 3B; Table S2). Therefore, based on genomic context and biological function, we propose to name Plu1493-like subfamily the group formed by the clades containing TseVs, Aave_0499, IdrD and PoNe (Fig. 3A, colors).

Multiple amino acid sequence alignments from each clade revealed the conserved residues characteristic of the PD-(D/E)xK superfamily (Fig. 3C), which comprise the aspartic acid (D), glutamic acid (E) and lysine (K) that are part of the catalytic site responsible for hydrolyzing phosphodiester bonds (Steczkiewicz *et al*., 2012). Using this information as a guide, substitution of the conserved aspartic acid for alanine in TseV2 and TseV3 (TseV_D282A_ and TseV3_D230A_) abrogated toxicity in *E. coli* (Fig. 3D). These results confirm that the enzymatic activity of the VRR-Nuc domain is essential for toxicity.

### TseV2 and TseV3 induce cell death via DNA double-strand breaks

We set out to determine whether TseV2 and TseV3 could cause DNA damage by analyzing the activation of the SOS response - a stress response mechanism induced by the activation of RecA (recombinase protein A) in response to DNA damage (Walker, 1996). *E. coli* harboring the reporter plasmid pSC101-P_*recA*_::GFP (Ronen *et al*, 2002), which carries the green fluorescent protein (GFP) under the control of the P_*recA*_ promoter, was co-transformed with either pBRA TseV2 or TseV3 and grown in AB media containing either 0.2% D-glucose or 0.2% L-arabinose (Fig. 4A and 4B). We observed an increase in GFP fluorescence when the expression of TseV2 or TseV3 was induced with L-arabinose, indicating the activation of the SOS response (Fig. 4A and 4B). To further assess the impact of TseV2 and TseV3 on bacterial chromosome stability, we used DAPI (4′,6-diamidino-2-phenylindole) to stain *E. coli* cells after inducing the expression of TseV2 or TseV3 for 1 h and evaluate nucleoid integrity by measuring the mean DAPI fluorescence per cell (Fig. 4C-F). Cells expressing TseV2 or TseV3 revealed smaller/degraded nucleoids and displayed reduced DAPI fluorescence (Fig. 4C-F), which indicates DNA degradation.

**Fig. 4.**
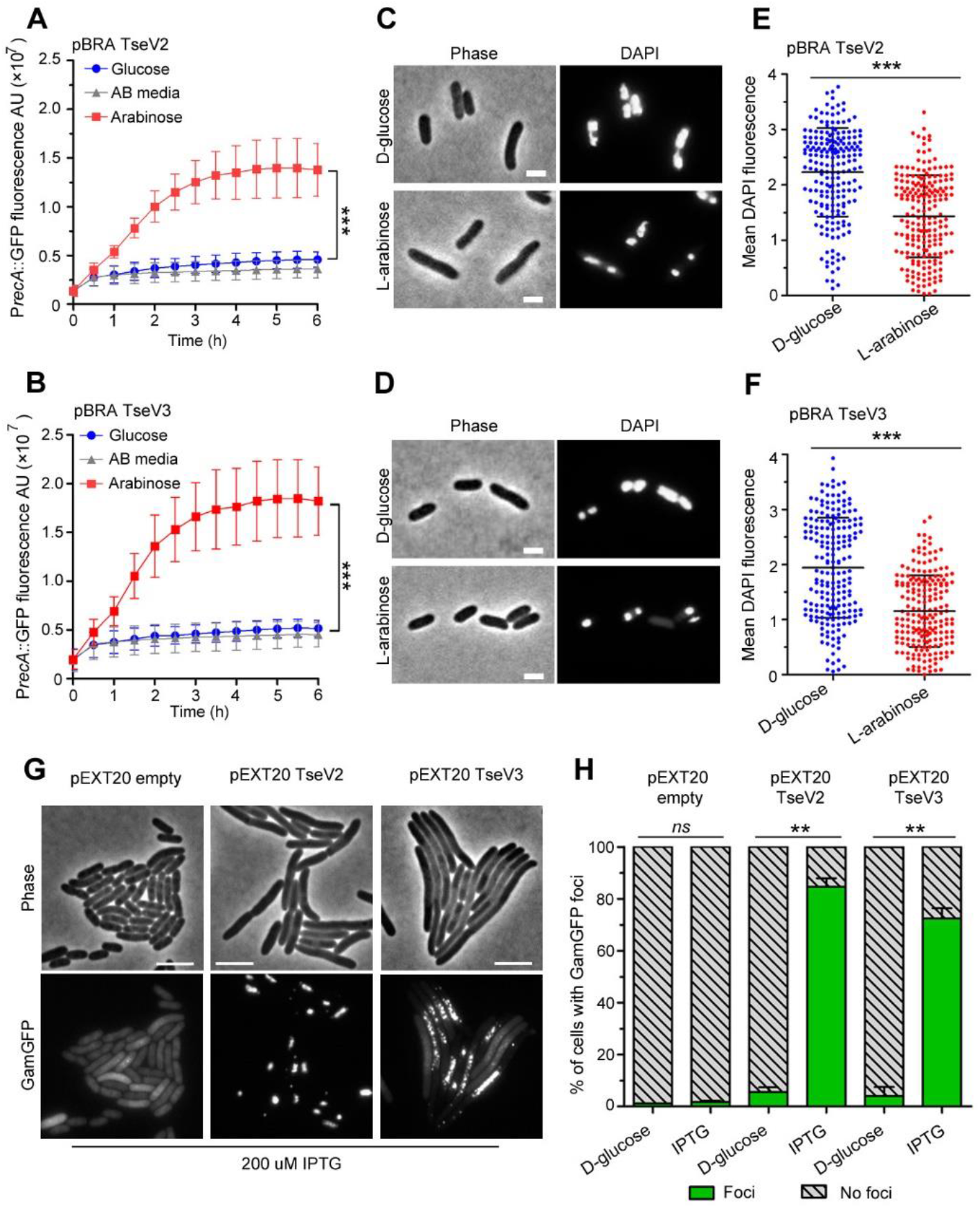
TseVs induce target cell death via DNA double-strand breaks. (A-B) Activation of the SOS response was analyzed using *E. coli* cells harboring the reporter plasmid pSC101-P_*recA*_::GFP and pBRA TseV2 (A) or pBRA TseV3 (B), which were grown in AB defined media with D-glucose or L-arabinose. Data is the mean ± SD of three independent experiments. ***p<0.001 (Student’s *t* test). (C-D) Bright-field and DAPI images of *E. coli* cells carrying pBRA TseV2 (C) or pBRA TseV3 (D) grown in the presence of D-glucose (repressed) or L-arabinose (induced). Results are representative images of three independent experiments. (E-F) Quantification of the mean DAPI fluorescence per cell of 200 cells. Data correspond to the mean ± SD of a representative experiment. Scale bar 2 μm. ***p < 0.001 (Student’s *t* test). (G) Representative bright-field and GFP images of *E. coli* co-expressing GamGFP and pEXT20 TseV2 or pEXT20 TseV3. Double-strand breaks appear as foci of GamGFP. Images are representatives of three independent experiments. Scale bar: 5 μm. (H) Quantification of the GamGFP foci shown in (G). Data are shown as the mean ± SD of the three independent experiments. **p < 0.01 (Student’s *t* test).

VRR-Nuc-containing enzymes have been shown to degrade 5’ flap single-strand DNA (Gwon *et al*., 2014) and Holliday junctions (a four-way junction in which two DNA double-strands are held together) (Pennell *et al*, 2014). Expression of TseV2 and TseV3 in *E. coli* followed by genomic and plasmid DNA extraction failed to reveal any significant difference in DNA integrity between induced and repressed conditions after visualization via electrophoresis in agarose gels (data not shown). To evaluate whether TseV2 and TseV3 could induce a small number of DNA double-strand breaks that were not detected by electrophoresis, we used the reporter strain *E. coli* SMR14354 encoding a chromosomal GFP fused to the Gam protein from bacteriophage Mu (GamGFP) under the control of the P_tet_ promoter (induced by tetracycline) (Shee *et al*, 2013). The Gam protein binds with high affinity and specificity to DNA double-strand ends, thus inducing the formation of GFP foci at specific sites (Shee *et al*., 2013). *E. coli* SMR14354 carrying an empty pEXT20 plasmid or encoding either TseV2 or TseV3 were grown with 0.2% D-glucose (repressed) or with 200 μM IPTG (induced) and examined by fluorescence microscopy (Fig. 4G and 4H). Cells carrying an empty plasmid revealed an even distribution of GamGFP in the cytoplasm, with only a few foci representing spontaneous double-strand breaks (Fig. 4G and 4H). Conversely, *E. coli* expressing either TseV2 or TseV3 revealed several intense GFP foci in more than 80% and 75% of cells, respectively (Fig. 4G and 4H). Interestingly, the expression of TseV2 leads to the formation of fewer intense foci per cell, whereas the expression of TseV3 induces the development of several less intense foci per cell (Fig. 4G and 4H), suggesting that TseV3 might cleave DNA at more sites than TseV2. Together, these results reveal that TseV2 and TseV3 cause target cell death via specific DNA double-strand breaks.

### TsiV3 interacts with the putative DNA-binding site of TseV3 to neutralize toxicity

To obtain information about the inhibitory mechanism of TsiV3, we decided to analyze whether this protein could directly interact with TseV3. Recombinant proteins were expressed, and the purified complex was analyzed using size exclusion chromatography coupled to multiple-angle light scattering (SEC-MALS) (Fig. S4). The MALS calculated average mass for the complex was 66.4 ± 3.3 kDa, which is close to the sum of the theoretical values of their monomers: 26.8 kDa and 37.4 kDa for 6xHis-TseV3 and TsiV3, respectively. SDS-PAGE analysis of the mixture confirmed the presence of 6xHis-TseV3 and TsiV3 (Fig. S4). These results reveal that TseV3 and TsiV form a 1:1 heterodimeric complex.

We were able to obtain crystals of the TseV3:TsiV3 complex, which belong to space group P2_1_1 and diffracted to a moderate resolution of 4 Å (Table S3). Matthews coefficient analysis indicated that two TseV3:TsiV3 complexes would be the most likely composition in the asymmetric crystal unit. We used AlphaFold (Jumper *et al*, 2021) models of TseV3_132-281_ and TsiV3_10-327_ for molecular replacement using Phaser (McCoy *et al*, 2007), which was able to place two copies of each monomer in the asymmetric unit with a final LLG (log-likelihood gain) of 486.87 and TFZ (translation function Z-score) of 12.4 - with both heterodimeric complexes adopting the same pose (our docked model is available using accession code ma-oyho8 at modelarchive.org). Therefore, the molecular replacement solution using the AlphaFold models most likely represents the correct relative orientation of the two subunits in the TseV3:TsiV3 complex (Fig. 5A). Given the relatively low resolution of the X-ray diffraction data, we chose not to refine these models against the processed dataset; however, our molecular replacement solution using the AlphaFold models was confirmed by identical placement using experimental PDB homologs taken from the DALI search described below - both TsiV and TsiT can be successfully utilized as search models for our experimental data, producing TFZ scores of 8.3 and 9.2, respectively. Attempts to co-fold the TseV3 and TsiV3 complex with AlphaFold did not result in the extensive interface we observe in our experimentally docked single models, thus confirming the requirement for data-derived docking.

**Fig. 5.**
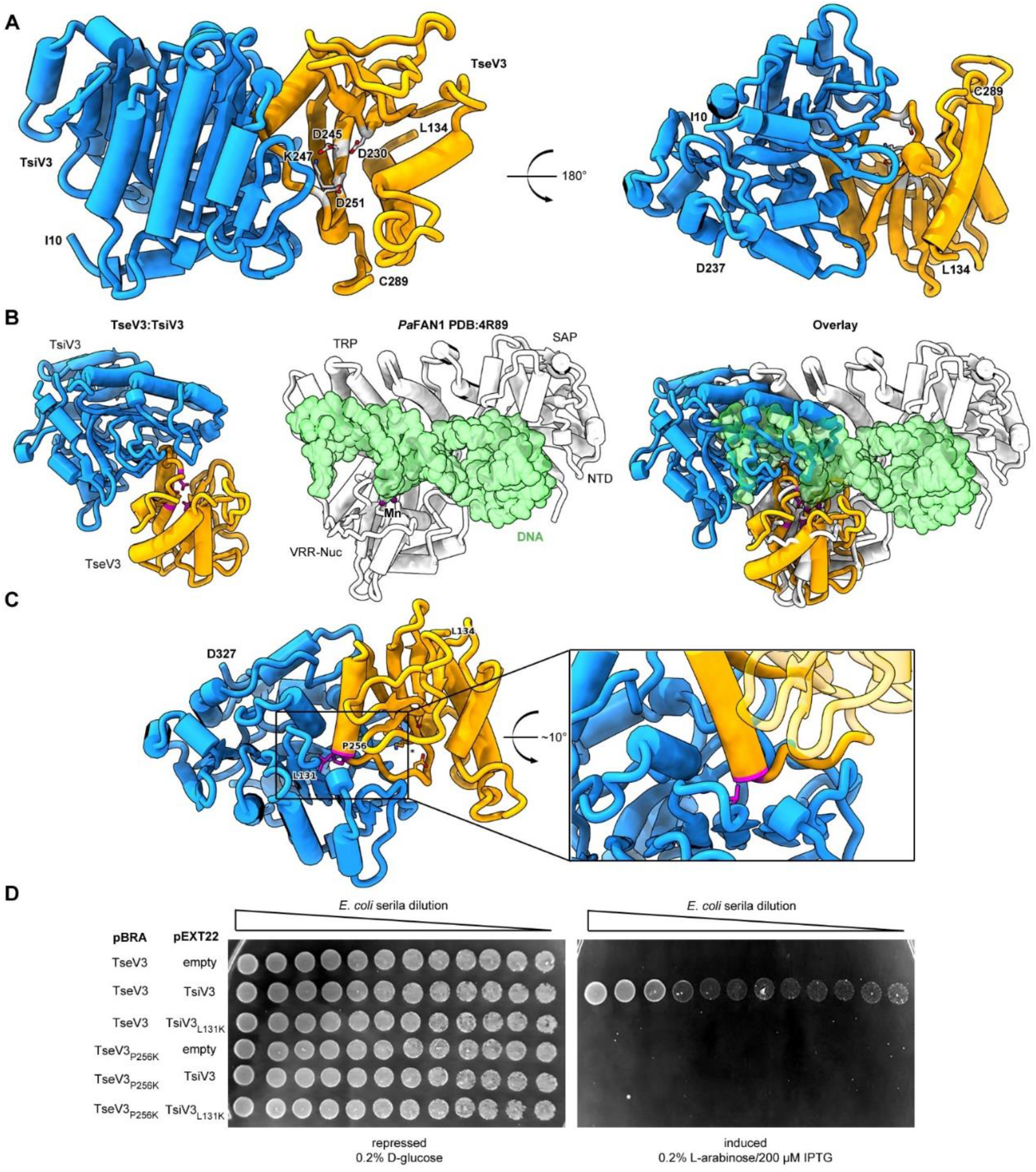
The effector-immunity complex reveals that TsiV3 blocks TseV3 substrate-binding site. (A) Constrained model of the TseV3:TsiV3 heterodimer with two different views: TsiV3 in blue (I_10_-D_327_) and TseV3 in orange (L_134_-C_289_). Models are labeled to assist interpretation. PD-(D/E)xK superfamily conserved residues of TseV3 (D_230_, D_245_, and K_247_) are shown in stick form and colored light grey, confirming that they converge to form a putative consensus active site. (B) Superimposition of the TseV3:TsiV3 coordinates with those of the *Pa*FAN1:DNA complex (PDB 4R89). *Pa*FAN1 protein in white, DNA duplex in green, and catalytic Mn^2+^ are depicted as purple spheres. The overlay (right) is presented in the same orientation as the individual complexes: TseV3:TsiV3 (left, catalytic residues in magenta) and *Pa*FAN1:DNA (middle). (C) Prediction of interface-compromising mutants in the TseV3:TsiV3 heterodimer. TsiV3 (blue) and TseV3 (orange) with putative active sites labeled with asterisk. Residues L_131_ of TsiV3 and P_256_ of TseV3 (both in stick form, magenta) form the closest point of contact in the heterodimer and are at the center of a hydrophobic-rich interface. (D) *E. coli* toxicity assay using cells carrying plasmids with wild-type or point mutations in TsiV3 (L_131_K) or TseV3 (P_256_K) as a potential means to destabilize the TseV3:TsiV3 complex interaction.

TsiV3 possesses a central β-sheet that is flanked by α-helices on one side and exposed on the other (Fig. 5A). This exposed β-sheet surface of TsiV3 binds to an α-β element of TseV3 that flanks its putative active site (composed of residues D_230_, D_245_, K_247_) (Fig. 5A). In this configuration, the TseV3-interacting α-helix, which corresponds to the α_2_ of the classical PD-(D/E)xK α_1_β_1_β_2_β_3_α_2_β_4_ core topology, projects its N-terminus towards the TsiV3 β-sheet (Fig. 5A).

Searches for structures similar to TseV3_132-281_ and TsiV3_10-327_ using the DALI server (Holm, 2020) revealed matches to proteins of related function: (a top Z-score of 33.5 for the immunity protein PA0821 PDB:7DRG and TsiV3; and a top Z-score of 3.8 between VRR-Nuc protein psNUC PDB:4QBL and TseV3). The modest RMSD (Root-Mean-Square Deviation) for the Cα positions in TseV3 and other VRR-Nuc enzymes indicates that TseV3 represents a variant of the VRR-Nuc family fold. Nevertheless, the PD-(D/E)xK consensus catalytic residues are identifiable as the modified sequence MD_230_IX_n_D_245_VK_247_ in TseV3. These residues are found in positions commensurate with active nucleases of the PD-(D/E)xK superfamily. Accordingly, superimposition of TseV3 with the well-characterized VRR-Nuc member *Pa*FAN1 (PDB 4R89) (Gwon *et al*., 2014) (clade KIAA1018 in Fig. 3A) matches residues D_230_, D_245_ and K_247_ of the former with residues D_507_, E_522_, and K_524_ of the latter (Fig. 5B).

*Pa*FAN1 (Gwon *et al*., 2014) is the bacterial homolog of the human FAN1 (MacKay *et al*, 2010), which is involved in the repair of DNA damage such as interstrand cross-links (ICLs) (Gwon *et al*., 2014; Jin *et al*, 2018). Human FAN1 is a structure-selective nuclease consisting of four domains: ubiquitin-binding zinc (UBZ); SAF-A/B, Acinus and PIAS (SAP); tetratricopeptide repeat (TPR); and VRR-Nuc (Iyer *et al*., 2006). Conversely, *Pa*FAN1 lacks the UBZ domain and contains an uncharacterized N-terminal domain (NTD), followed by SAP, TPR and VRR-Nuc (Iyer *et al*., 2006; Gwon *et al*., 2014). The structure of *Pa*FAN1 has been solved in complex with a 5’ flap DNA substrate (PDB 4R89) (Fig. 5B, middle). As the catalytic residues of TseV3 align with those of *Pa*FAN1, we used the structure of the latter as a guide to analyze the likely mechanism by which TsiV3 may neutralize TseV3 activity. Comparison between the *Pa*FAN1:DNA and TseV3:TsiV3 complexes reveals that the path of the DNA substrate is potentially incompatible with the presence of TsiV3 (Fig. 5B). Hence, assuming the mode of substrate recognition is similar between *Pa*FAN1 and TseV3, this result suggests that the binding of TsiV3 sterically blocks the toxin active site. In our model of the complex, TsiV3 occlusion of the TseV3 active site would be enabled by TsiV3 loop, which projects into the putative TseV3 DNA-binding pocket (Fig. 5A, B).

To validate the accuracy of our TseV3:TsiV3 structural model, we designed point mutations to disrupt the interaction surface between the two proteins without perturbing the active site or its ability to bind DNA. Thus, residue P256 at the N-terminus of α_2_ of TseV3 was replaced by lysine (TseV3_P256K_), and residue L131 in the central β-sheet of TsiV3 was also replaced by lysine (TsiV3_L131K_) (Fig. 5C). Native and mutated versions of the effector and immunity protein were co-transformed in *E. coli* to analyze toxicity (Fig. 5D). Results revealed that TsiV3_L131K_ was unable to neutralize TseV3 toxicity. In addition, the mutated TseV3_P256K_ maintained its enzymatic activity, displaying toxicity in *E. coli*; however, this mutant was not neutralized by co-expression with the native immunity protein (TsiV3) (Fig. 5D). Together, these results reinforce the accuracy of our model, which is comprised of both experimental constraints and theoretical model components.

## Discussion

Bacterial antagonistic strategies targeting nucleic acids are very effective as these components are critical for life. In this study, we characterized a group of effectors containing the VRR-Nuc domain. This domain has not previously been reported to be used in biological conflicts (Zhang *et al*., 2012), but recently was suggested to work as a T6SS effector due to its localization next to a PAAR protein in *P. aeruginosa* (Wang *et al*., 2021) – for consistency we have decided to keep the name TseV for this group of effectors. Proteins containing the VRR-Nuc domain comprise a family (Iyer *et al*., 2006) belonging to the PD-(D/E)xK superfamily, which contain a conserved enzymatic core composed by α_1_β_1_β_2_β_3_α_2_β_4_ (Steczkiewicz *et al*, 2012). The conserved catalytic residues (D, E, K) are located in the central β_2_β_3_-sheet, while the α_1_-helix is associated with the formation of the active site and α_2_-helix with substrate binding (Steczkiewicz *et al*., 2012). Curiously, *S. bongori* encodes four TseV homologs: TseV2 and TseV3 are toxic in *E. coli*, whereas TseV1 and TseV4 are not toxic (Fig. 2). Based on what is known about the catalytic mechanism of PD-(D/E)xK nucleases, we hypothesize that the lack of α_2_- and α_1_-helix in TseV1 and TseV4, respectively, might explain the lack of toxicity (Fig. S5). Another curiosity is the presence of two homologs of the DUF3396 immunity genes downstream of both TseV2 and TseV1 (Fig. 2A). Such genomic organization is also conserved in other bacterial species like *Photorhabdus thracensis* (VY86_01065, VY86_01040), *Photorhabdus asymbiotica* (PAU_03539, PAU_03660) (Enterobacterales), *Marinobacter nauticus* (MARHY2492) (Pseudomonadales) and *Herbaspirillum huttiense* (E2K99_00955) (Burkholderiales). The fact that only one immunity protein (TsiV2.1) can neutralize the effector (TseV2) makes us wonder about the role of the additional immunity protein gene - and why such genomic context is conserved. One possibility is that the extra immunity protein could regulate the effector at the transcriptional level as has been reported for the immunity protein TsiTBg known to regulate a different PD-(D/E)xK effector (TseTBg) (Yadav *et al*., 2021).

The complexity of the PD-(D/E)xK superfamily and the rapid evolution of polymorphic toxins makes it difficult to categorize antibacterial effectors belonging to this group. However, our phylogenetic analysis was able to confidently group VRR-Nuc-containing effectors in one clade (TseV) and show that this group is different from the clades formed by the homologs of additional T6SS effectors (Aave_0499, IdrD, PoNe) (Fig. 3). Although proteins belonging to clades Aave_0499, IdrD, PoNe are not recognized by the Pfam model of VRR-Nuc, these proteins share similar genetic architectures concerning domain fusions and gene vicinity (Fig. 3B); therefore, we decided to call this larger group Plu1493-like to respect the original nomenclature proposed by Iyer et al. (2006).

The enzymatic activity of proteins belonging to the PD-(D/E)xK superfamily is quite diverse, but we were able to narrow down the possibilities and reveal that TseV effectors intoxicate target cells by inducing DNA double-strand breaks. Despite killing cells by the same mechanism, *E. coli* intoxicated by TseV2 and TseV3 display slight differences in phenotype. Due to the difference in GamGFP phenotype (Fig. 4G) and the number of resistant colonies obtained after ectopic expression of TseV2 and TseV3 in *E. coli* (Fig. S1), we hypothesize that the former cleaves DNA at fewer sites compared to the latter. It remains to be defined whether these effectors target a specific nucleotide sequence or recognize some conformational DNA structure. Unfortunately, whole genome sequencing of 20 resistant *E. coli* clones expressing the toxin (Fig. S1) failed to uncover any evident mutation that could be attributed to the resistance phenotype (data not shown).

T6SSs effector-immunity complexes are related to type II toxin-antitoxin (TA) systems, which play several roles in bacterial physiology ranging from genomic stabilization and abortive phage infection to stress modulation and antibiotic persistence (Fraikin *et al*, 2020). Most T6SS immunity proteins described to date bind to effectors to regulate their enzymatic activity (Benz *et al*, 2012; Benz *et al*., 2013; Dong *et al*, 2013; Li *et al*, 2013; Lu *et al*, 2014; Robb *et al*, 2016). An exception is Tri1 (type VI secretion ADP-ribosyltransferase immunity 1) from *Serratia proteamaculans*, which exhibits two modes of inhibition: active site occlusion and enzymatic removal of a post-translational modification (Ting *et al*, 2018). The neutralization mechanism of TsiT, which counteracts the PD-(D/E)xK effector TseT from *P. aeruginosa*, was also proposed to be different: TsiT interferes with the effector oligomerization state and hinders its nuclease activity (Wen *et al*, 2021). Our structural model of the TseV3:TsiV3 complex revealed that TsiV3 β-sheet binds to the α_2_-helix of TseV3, which is involved in DNA binding in other PD-(D/E)xK members (Steczkiewicz *et al*., 2012). In addition, the superposition of the TseV3:TsiV3 complex with the structure of *Pa*FAN1 bound to DNA confirms the hypothesis that TsiV3 likely occludes the substrate-binding site of TseV3.

It is estimated that *E. coli* and *Salmonella* diverged millions of years ago (Fookes *et al*., 2011). *Salmonella*-specific functions are encoded by genes located in prophages and specific SPIs. Several characteristics of *S. bongori* suggest that this species may lie somewhere between *E. coli* and *S. enterica* during evolution (Christensen *et al*, 1998; Fookes *et al*., 2011). Analyzing the synteny between *S*. Typhimurium 14028s and *S. bongori* NCTC 12419, we observed that the SPI-2 T3SS of *S. enterica* is localized in the equivalent genomic loci of the SPI-22 T6SS of *S. bongori* (Fig. S6). The SPI-2 T3SS of *S. enterica* is well characterized for its importance during the intracellular stage of the infection cycle, allowing bacteria to manipulate host cellular functions and replicate (Hensel *et al*, 1995; Jennings *et al*, 2017). Whether the SPI-22 T6SS of *S. bongori* also works against eukaryotic cells remains to be established.

The genome of *S. bongori* NCTC 12419 encodes a large repertoire of orphan Hcp, VgrG and PAAR-like proteins (Fig. 1B) that are responsible for diversifying the array of effectors secreted by the SPI-22 T6SS. Conversely, the genome of *S*. Typhimurium 14028s that encodes a phylogenetically unrelated SPI-6 T6SS involved in competition with species of the gut microbiota (Sana *et al*., 2016; Sibinelli-Sousa *et al*., 2020) displays a restricted repertoire of orphan T6SS genes composed of only two Hcps (STM14_3785 and STM14_5414) (Blondel *et al*., 2009). These observations suggest that the former bacterium may use its SPI-22 T6SS to target a greater variety of competitors, while the latter uses its SPI-6 T6SS to target a restricted number of species.

Here we add to the known diversity of antibacterial weapons, placing the VRR-Nuc family within the remit of T6SS effectors. Knowledge about the phylogeny and mechanism of action of this group of effectors will be important in interpreting its usage in other bacterial species, including the requirements of neutralization by very specific immunity pairings.

## Material and Methods

### Bacterial strains and growth conditions

A list of bacterial strains used in this work can be found in Table S4. Strains were grown at 37°C in Lysogeny Broth (10 g/L tryptone, 10 g/L NaCl, 5 g/L yeast extract) under agitation. Cultures were supplemented with antibiotics in the following concentration when necessary: 50 μg/mL kanamycin, 100 μg/mL ampicillin, and 50 μg/mL streptomycin.

### Cloning and mutagenesis

Putative effectors SBG_1828, SBG_1841, SBG_2718 and SBG_2723 were amplified by PCR and cloned into pBRA vector under the control of P_BAD_ promoter (Souza *et al*, 2015). Immunity proteins SBG_1828, SBG_1842, SBG_2719, SBG_2720, SBG_2724 and SBG_2725 were cloned into pEXT22 under the control of P_TAC_ promoter (Dykxhoorn *et al*, 1996). TseV2 and TseV3 were cloned in the pEX20 vector under the control of P_TAC_ promoter (Dykxhoorn *et al*., 1996) for GamGFP assays. For complementation, SBG_1238 (TssB), SBG_1842 (TsiV3) and SBG_2724 (TsiV2.1) were cloned into pFPV25.1 by replacing the GFP mut3 coding region for the genes of interest (Valdivia & Falkow, 1996). Point mutations were created using QuikChange II XL Site-Directed Mutagenesis Kit (Agilent Technologies) and pBRA TseV2 and pBRA TseV3 plasmids as templates. S. *bongori* mutant strains were constructed by λ-Red recombination engineering using a one-step inactivation procedure (Datsenko & Wanner, 2000). All constructs were confirmed by sequencing.

### Interbacterial competition assay

Bacterial competition assays were performed using *S. bongori* (WT, Δ*tssB*, Δ*tseV2/tsiV2*.*1/tsiV2*.*2* and Δ*tseV3/tsiV3*) as attackers, and *E. coli* K-12 W3110 carrying pFPV25.1 Amp^R^ as prey. Overnight cultures of the attacker and prey cells were sub-cultured in LB (1:30) until reaching OD_600ηm_ 1.6, then adjusted to OD_600ηm_ 0.4 and mixed in a 10:1 ratio (attacker:prey), 5 μL of the mixture were spotted onto 0.22 μm nitrocellulose membranes (1 × 1 cm) and incubated on LB agar (1.5%) at 37°C for the indicated periods. Membranes containing the bacterial mixture were placed on 1.5 mL tubes containing 1 mL of LB, homogenized by vortex, serially diluted, and plated on selective plates containing antibiotics. The prey recovery rate was calculated by dividing the CFU (colony forming units) counts of the output by the CFU of the input.

### *E. coli* toxicity assays

Overnight cultures of *E. coli* DH5α (LB with 0.2% glucose) carrying effectors (in pBRA) and immunity proteins (in pEXT22) were adjusted to OD_600ηm_ 1, serially diluted in LB (1:4) and 5 μL were spotted onto LB agar plates containing either 0.2% D-glucose or 0.2% L-arabinose plus 200 μM IPTG – both supplemented with streptomycin and kanamycin - and incubated at 37°C for 20 h. For growth curves, overnight cultures of *E. coli* carrying pBRA TseV2 or TseV3 were inoculated in LB (1:50) with 0.2% D-glucose and grown at 37°C (180 rpm) for 1.5 h. Next, media was replaced with either fresh warm LB containing 0.2% D-glucose or 0.2% L-arabinose.

### Microscopy Time-lapse studies

For time-lapse microscopy, LB agar (1.5%) pads were prepared by cutting a rectangular piece out of a double-sided adhesive tape, which was taped onto a microscopy slide as described previously (Bayer-Santos *et al*, 2019). *E. coli* DH5α harboring pBRA TseV2 or TseV3 were sub-cultured in LB (1:50) with 0.2% D-glucose until reaching OD_600ηm_ 0.4-0.6 and adjusted to OD_600ηm_ 1.0. Cultures were spotted onto LB agar pads supplemented either with 0.2% D-glucose or 0.2% L-arabinose plus antibiotics. Images were acquired every 15 min for 16 h using a Leica DMi-8 epifluorescent microscope fitted with a DFC365 FX camera (Leica) and Plan-Apochromat 63x oil objective (HC PL APO 63x/1.4 Oil ph3 objective Leica). Images were analyzed using FIJI software (Schindelin *et al*, 2012).

### Bioinformatic analysis

Iterative profile searches using JackHMMER (Eddy, 2011) with a cutoff e-value of 10^−6^ and a maximum of four iterations were performed to search a non-redundant (nr) protein database from the National Center for Biotechnology Information (NCBI) (Sayers *et al*, 2019). Similarity-based clustering of proteins was carried out using MMseqs software (Steinegger & Söding, 2017). Sequence alignments were produced with MAFFT local-pair algorithm (Katoh & Standley, 2013), and non-informative columns were removed with trimAl software (Capella-Gutiérrez *et al*, 2009). Approximately-maximum-likelihood phylogenetic trees were built using FastTree 2 (Price *et al*, 2010). Sequence logos were generated using Jalview (Waterhouse *et al*, 2009). HMM models were produced for each sequence alignment and compared against each other with the HH-suit package (Steinegger *et al*, 2019). Proteins were annotated using the HHMER package (Eddy, 2011) or HHPRED software (Söding *et al*, 2005) and Pfam (Bateman *et al*, 2004), PDB (Berman *et al*, 2007) or Scope (Fox *et al*, 2014) databases. An in-house Python script was used to collect the gene neighborhoods based on information downloaded from the complete genomes and nucleotide sections of the NCBI database (Sayers *et al*., 2019).

TseV1-4 sequence alignments were produced with MAFFT local-pair algorithm (Katoh & Standley, 2013) and analyzed in AilView (Larsson, 2014) to separate the regions of interest. Sequence logos were produced using the Jalview (Waterhouse *et al*., 2009). Protein structure predictions were performed with ColabFold (Mirdita *et al*, 2021) and AlphaFold (Jumper *et al*., 2021), and visualization was performed using Pymol (DeLano, 2002).

The genome of *S. bongori* NCTC 12419 and *S*. Typhimurium NCTC 14028s were retrieved from the NCBI database and aligned with BLASTn (Camacho *et al*, 2009). The alignment was analyzed using Artemis Comparison Tool (ATC) (Carver *et al*, 2005). The genome map was constructed using the ATC plug-in DNAPlotter (Carver *et al*, 2009).

### SOS response assays

Overnight cultures of *E. coli* DH5α harboring the reporter plasmid pSC101-P_recA_::GFP (Ronen *et al*., 2002) and pBRA TseV2 or TseV3 were sub-cultured (1:50) in LB with 0.2% D-glucose and grown at 37ºC until OD_600ηm_ 0.4-0.6. Bacteria were harvested and resuspended in AB defined media (0.2% (NH_4_)_2_SO_4_, 0.6% Na_2_HPO_4_, 0.3% KH_2_PO_4_, 0.3% NaCl, 0.1 mM CaCl_2_, 1 mM MgCl_2_, 3 μM FeCl_3_) supplemented with 0.2% sucrose, 0.2% casamino acids, 10 μg/mL thiamine and 25 μg/mL uracil (Bayer-Santos *et al*., 2019). Cells (OD_600ηm_ 1.0) were placed in a black 96 well plate with clear bottom (Costar) with 0.2% D-glucose or 0.2% L-arabinose to a final volume of 200 μL. GFP fluorescence was monitored in a plate reader SpectraMax Paradigm Molecular Devices for 6 h at 30°C.

### DAPI staining

*E. coli* DH5α carrying pBRA TseV2 and TseV3 were sub-cultured in LB with 0.2% D-glucose (1:50) and grown at 37°C (180 rpm) until OD_600ηm_ 0.4-0.6. Cells were harvested and resuspended in new media with 0.2% D-glucose or 0.2% L-arabinose and growth for an additional 1 h. Bacteria were fixed with 4% paraformaldehyde for 15 min on ice, washed in phosphate buffer saline (PBS) and stained with DAPI (3 μg/mL) for 15 min at room temperature. Samples were washed once with PBS before transferring 1 µL of each culture to a 1.5% PBS-agarose pad for visualization. Images were acquired in Leica DMi-8 epifluorescent microscope fitted with a DFC365 FX camera (Leica) and Plan-Apochromat 63x and 100x oil objectives (HC PL APO 63x and 100x/1.4 Oil ph3 objectives Leica). Images were analyzed using FIJI software (Schindelin *et al*., 2012). To assess DNA integrity, the mean pixel fluorescence per cell was manually measured from 200 bacteria from different fields from each experiment. The cell area was determined using the bright field, and the mean pixel fluorescence per cell was measured in the DAPI channel subtracting the background.

### DNA double-strand break microscopy

*E. coli* SMR14354 containing a chromosomal GamGFP under the control of P_tet_ promotor (Shee *et al*., 2013) and harboring an empty pEXT20 or encoding TseV2 or TseV3 were sub-cultured in LB (1:100) with 0.2% D-glucose grown for 1.5 h at 37°C (180 rpm) before the induction of GamGFP with 50 ηg/mL tetracycline for 2 h. Bacteria were resuspended in new media with either 0.2% D-glucose or 200 μM IPTG and grown for 1 h. One microliter of each culture was spotted onto a 1.5% LB-agarose pad. Images were acquired in a Leica DMi-8 epifluorescent microscope fitted with a DFC365 FX camera (Leica) and Plan-Apochromat 100x oil objective (HC PL APO 100x/1.4 Oil ph3 objective Leica). Images were analyzed using FIJI software (Schindelin *et al*., 2012). At least 400 bacteria from each experiment were quantified.

### Protein expression and purification

*E. coli* SHuffle cells carrying pRSFDuet 6xHis-TseV3-TsiV3 were grown in LB supplemented with kanamycin (30°C, 180 rpm) until OD_600ηm_ 0.4-0.6. Expression was induced with 0.5 mM IPTG followed by incubation at 16°C for 16 h. Cells were harvested via centrifugation at 9000 *g* for 15 min, and pellets were resuspended in buffer A (50 mM Tris-HCl pH 7.5, 200 mM NaCl, 5 mM imidazole) and lysed at 4°C using an Avestin EmulsiFlex-C3 homogenizer. The lysate was collected and centrifuged (48000 *g*) for 1 h at 4 °C. The supernatant was loaded onto a 5 ml HisTrap HP cobalt column (GE Healthcare) equilibrated in buffer A. The column was washed with 10 column volumes (CV) of buffer A before running an elution gradient of 0-50% buffer B (50 mM Tris-HCl pH 7.5, 200 mM NaCl, 500 mM imidazole) over 10 CV, followed by a final 10 CV wash with 100% buffer B. The presence of TseV3-TsiV3 was confirmed by SDS-PAGE of eluted fractions. TseV3-TsiV3 was concentrated using a Vivaspin® spin-concentrator and further purified by size-exclusion chromatography on a Superdex 200 26/60 column (GE Healthcare) equilibrated in 50 mM Tris-HCl pH 7.5, 150 mM NaCl.

SEC-MALS analyses were used to determine the molar mass of the TseV3-TsiV3 complex (concentration 3.2 mg/mL). Protein samples (400 μL injection volume) were separated using a Superdex 200 10/300 column (GE Healthcare) equilibrated with buffer (50 mM Tris-HCL pH 7.5, 20 mM NaCl) coupled to a miniDAWN TREOS multi-angle light scattering system and an Optilab rEX refractive index detector. Data analysis was performed using the Astra Software package version 7.1 (Wyatt TechnologyCorp). Molecular mass was calculated assuming a refractive index increment dn/dc = 0.185 mL/g (Wen *et al*, 1996). Fractions were analyzed in SDS-PAGE to confirm protein molecular weight.

### Crystallography and structure determination

TseV3-TsiV3 was concentrated to 18 mg/mL and crystalized in 0.1 M HEPES pH 7.5 and 30 % v/v PEG Smear Low (12.5% v/v PEG 400, 12.5% v/v PEG 500, monomethylether, 12.5% v/v PEG 600, 12.5% v/v PEG 1000). The crystals were cryoprotected in the mother liquor supplemented with 20% ethylene glycol and subsequently cryo-cooled in liquid nitrogen. X-ray diffraction data were collected at Diamond Light Source on beamline i04, and initial data processing was performed using the xia2-dials pipeline (Winter, 2010; Winter *et al*, 2018). The data were phased by molecular replacement in Phaser (McCoy *et al*., 2007) using AlphaFold (Jumper *et al*., 2021) models of TseV3_134-289_ and TsiV3_10-327_, which were trimmed to include only the high-confidence regions and omit the N-terminal DUF4150 domain of TseV3.

### Quantification and statistical analyses

Statistical test, number of events, mean values and standard deviations are reported in each figure legend accordingly. Statistical analyses were performed using GraphPad Prism5 software and significance is determined by the value of p < 0.05.

## Acknowledgements

We are grateful to Cristiane Rodrigues Guzzo for sharing reagents and equipment, and Alexandre Bruni Cardoso for allowing access to the fluorescence microscope. We thank members of the LEEP and EBS groups for scientific discussions. Crystallography analyses were performed at Diamond Light Source. This work was supported by São Paulo Research Foundation (FAPESP) grants to RFS (2016/09047-8), CSF (2017/17303-7) and EBS (2017/02178-2). AL is supported by Welcome Trust (209437/Z/17/Z). FAPESP fellowships were awarded to JTH (2018/25316-4), DESL (2019/22715-8), GGN (2021/03400-6), EEL (2019/12234-2), GSC (2020/15389-4), and EBS (2018/04553-8). LM is supported by a MIBTP studentship.

## Author contributions

JTH, DESL and EBS outlined the study. JTH, DESL, EEL, GSC, and EBS performed experiments and analyzed data. LM and AL performed protein crystallography and analyzed data. GGN and RFS contributed with bioinformatic analyses. JTH, DESL, GGN, CSF, RFdS, RSG, AL and EBS contributed to the scientific discussions. JTH and EBS wrote the manuscript with input from other authors. All authors revised and approved the manuscript.

## Supplementary Figures

**Fig. S1.**
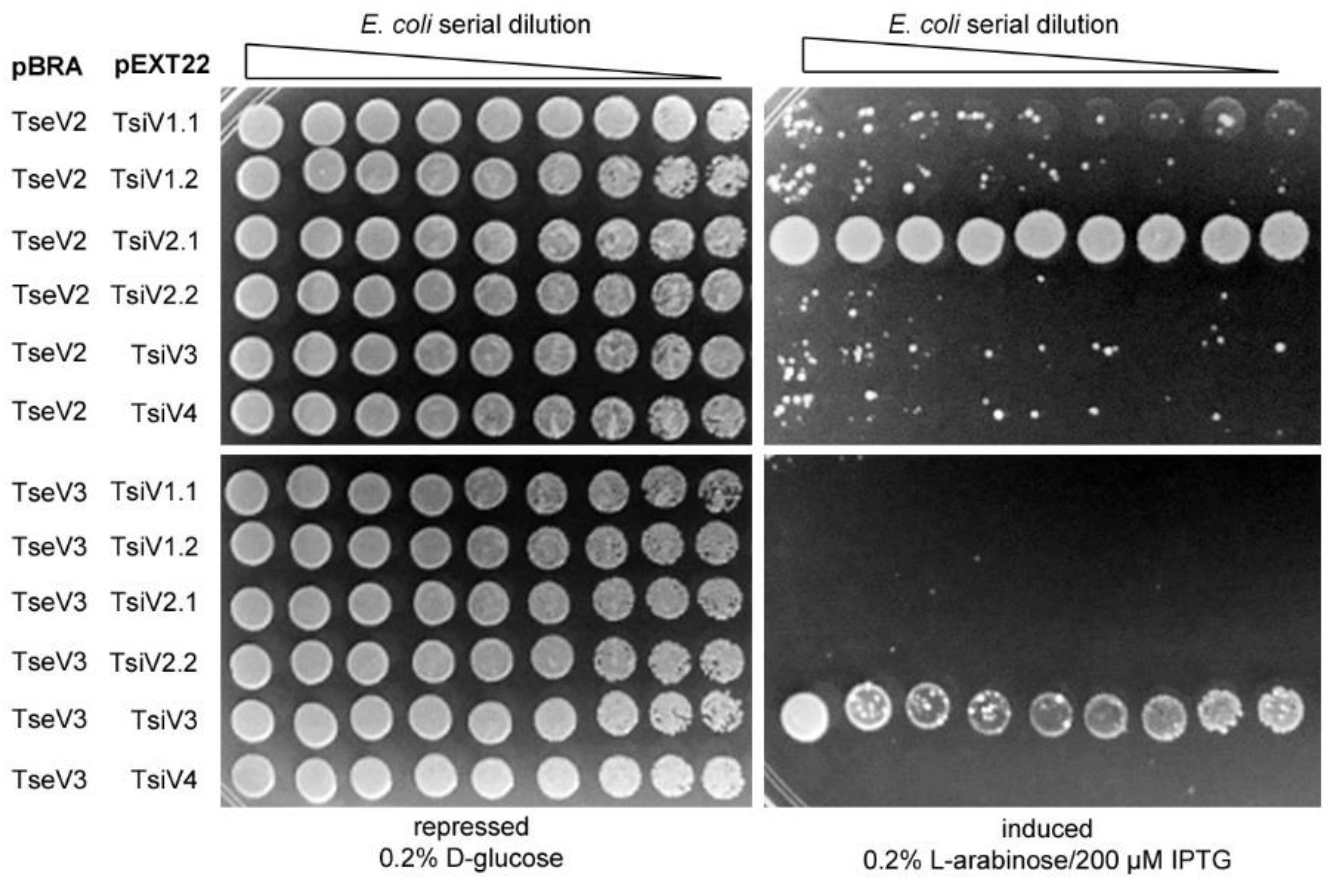
Toxicity assay in *E. coli* co-transformed with pBRA TseV2 or pBRA TseV3 and the six different immunity proteins. Only one specific immunity could abrogate the toxic effect. Images are representative of three independent experiments.

**Fig. S2.**
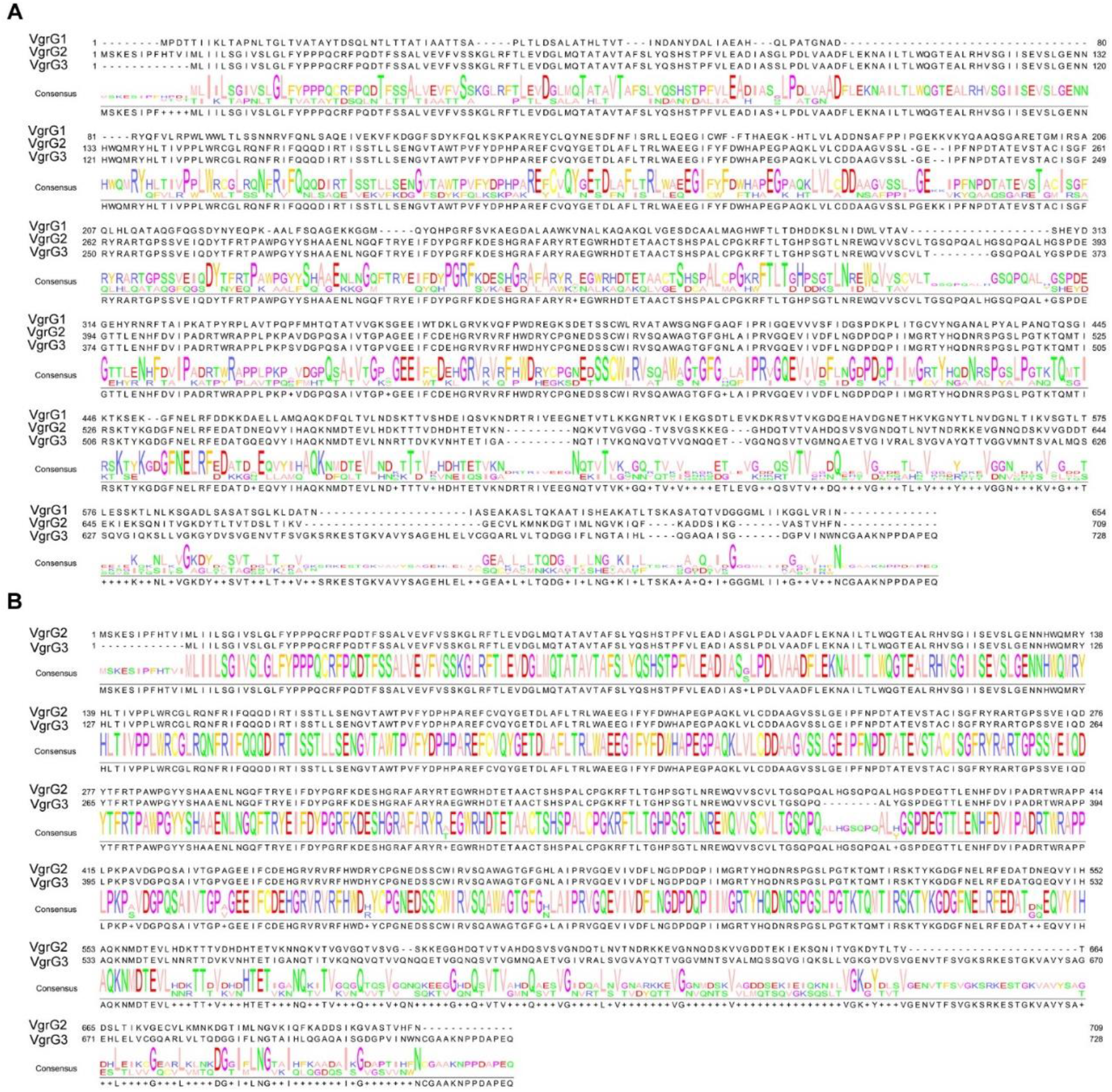
(A) Amino acid sequence alignment of VgrG1, VgrG2 and VgrG3. (B) Amino acid sequence alignment of VgrG2 and VgrG3. Amino acids are color-coded according to their properties.

**Fig. S3.**
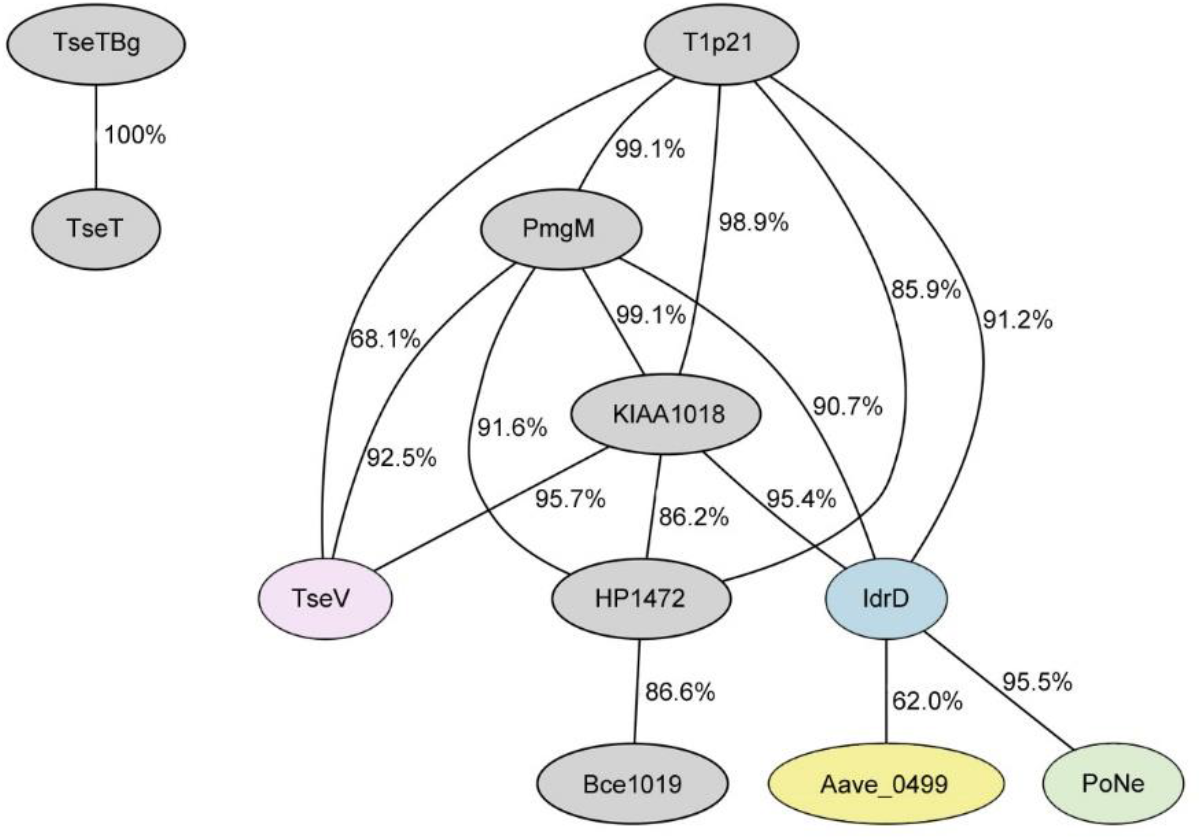
Comparison of the HMM (Hidden Markov Model) from each clade shown in Fig. 3A. All clades present enough similarity to be clustered together, while the homologs of *P. aeruginosa* TseT and *B. gladioli* TseTBg differ from the other models.

**Fig. S4.**
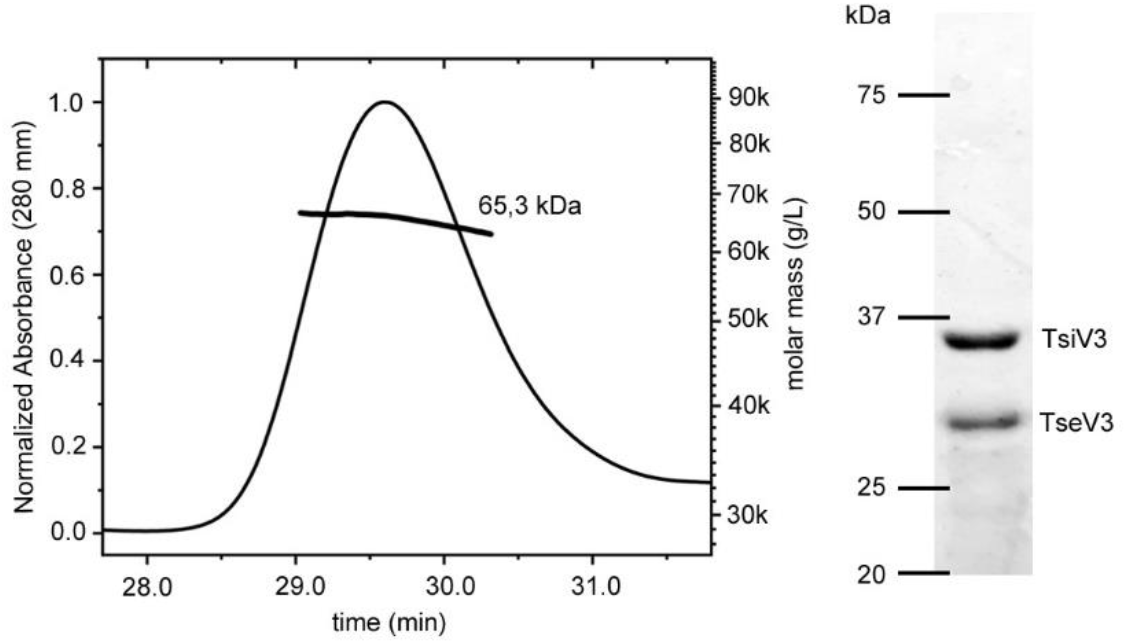
SEC-MALS analysis shows the formation of a stable complex between TseV3:TsiV3. The line corresponds to the calculated molecular mass. Right panel: SDS-PAGE showing the apparent molecular mass of proteins eluted from SEC-MALS peak.

**Fig. S5.**
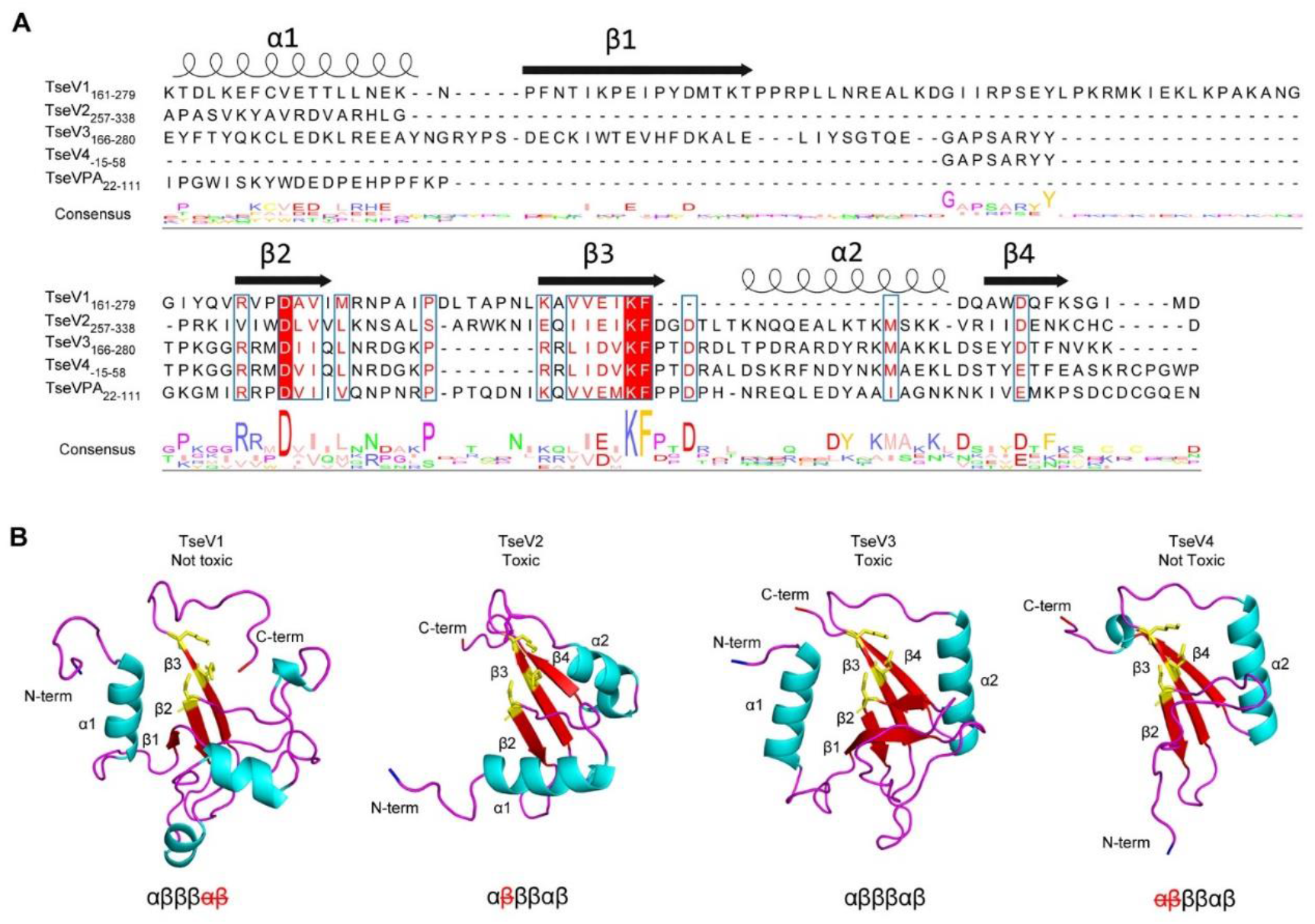
(A) Manual amino acid sequence alignment of TseV1-4 and *P. aeruginosa* TseV (PA0822) based on secondary structures. The secondary structures are indicated above the alignments with α-helixes represented by spirals and β-sheets by arrows. The conserved catalytic residues are highlighted in red with the logo underneath the alignments. TseV4 contains another start codon located upstream of the annotated one. (B) TseV1-4 structures predicted by the AlphaFold (Jumper *et al*., 2021). Underneath is the conserved PD-(D/E)xK enzymatic core with the absent structures marked in dashed red.

**Fig. S6.**
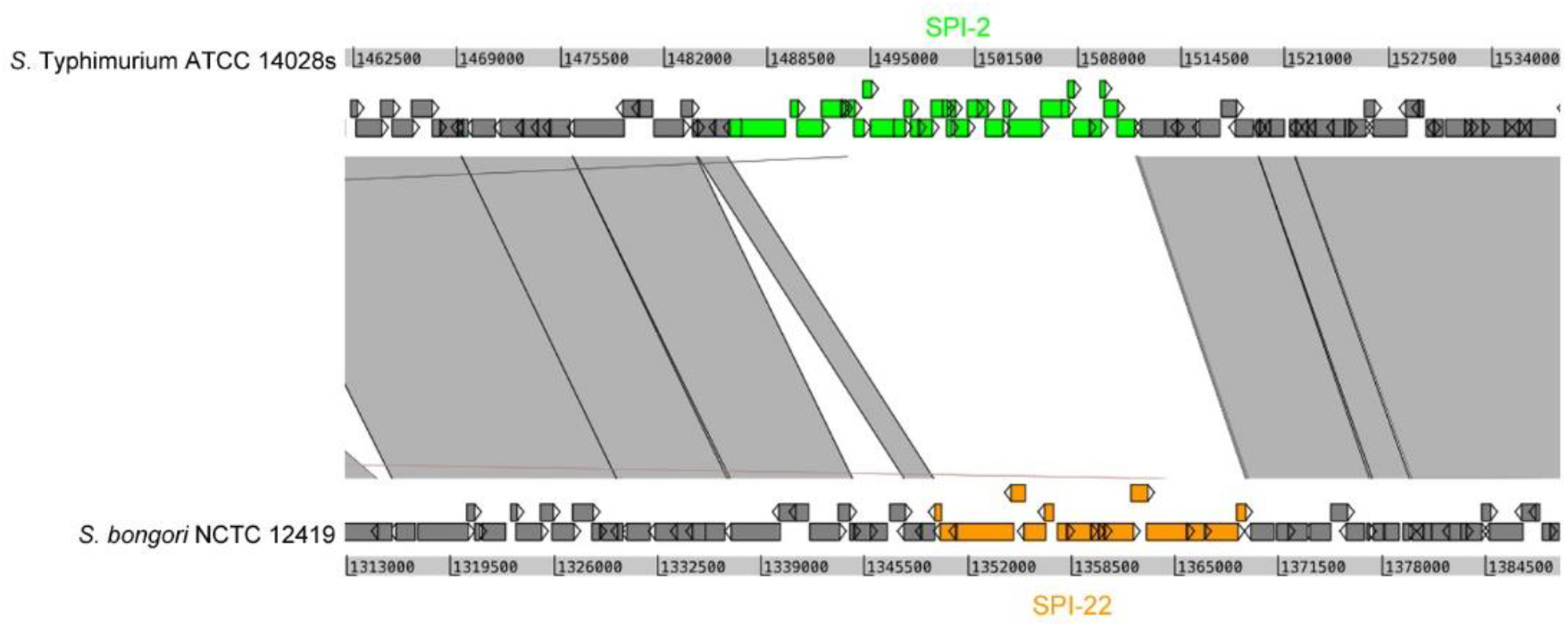
Genomic alignment comparing *S*. Typhimurium and *S. bongori*. The regions encoding *S*. Typhimurium SPI-2 T3SS and *S. bongori* SPI-22 T6SS are in focus.

## Supplementary Legends

**Table S1A-J**. List of all homologs collected by JackHMMER searches and used to build the phylogenetic tree shown in Fig. 3A.

**Table S2A-J**. List of genes surrounding representative sequences from each VRR-Nuc subfamily.

**Table S3**. Crystallographic statistics of the TseV3:TsiV3 complex.

**Table S4**. Strains, plasmids and primers used in the study.

**Movie S1**. Time-lapse microscopy of *E. coli* harboring pBRA TseV2 growing in media supplemented with 0.2% D-glucose. Timestamp in hh:mm. Scale bar: 5 μm. Arrows indicate selected bacteria shown in Fig. 2D.

**Movie S2**. Time-lapse microscopy of *E. coli* harboring pBRA TseV2 growing in media supplemented with 0.2% L-arabinose. Timestamp in hh:mm. Scale bar: 5 μm. Arrows indicate selected bacteria shown in Fig. 2D.

**Movie S3**. Time-lapse microscopy of *E. coli* harboring pBRA TseV3 growing in media supplemented with 0.2% D-glucose. Timestamp in hh:mm. Scale bar: 5 μm. Arrows indicate selected bacteria shown in Fig. 2D.

**Movie S4**. Time-lapse microscopy of *E. coli* harboring pBRA TseV3 growing in media supplemented with 0.2% L-arabinose. Timestamp in hh:mm. Scale bar: 5 μm. Arrows indicate selected bacteria shown in Fig. 2D.

